# Bovine leukemia virus-derived long-noncoding RNA, AS1-S, binds to bovine hnRNPM and alters interaction properties between hnRNPM and host mRNAs

**DOI:** 10.1101/2023.02.27.530377

**Authors:** Kiyohiko Andoh, Asami Nishimori, Yuichi Matsuura

**Author notes:** Address correspondence to Kiyohiko Andoh. E-mail addresses: Kiyohiko Andoh, Asami Nishimori, Yuichi Matsuura.

## Abstract

Viruses utilize several strategies to develop latent infection and evade host immune responses. Long non-coding RNA (lncRNA), a class of non-protein encoding RNA that regulates various cellular functions by interacting with RNA binding proteins, is a key factor for viral latency because of its lack of antigenicity. Bovine leukemia virus (BLV), which belongs to the family Retroviridae, encodes the BLV-derived lncRNA AS1-S, which is a major transcript expressed in latently infected cells. We herein identified bovine hnRNPM, an RNA-binding protein located in the nucleus, as the binding partner for AS1-S using an RNA-protein pull-down assay. The pull-down assay using recombinant hnRNPM mutants showed that RNA recognition motif 1 and 2, located in the N-terminal region of bovine hnRNPM, are responsible for binding AS1-S. Furthermore, an RNA immunoprecipitation assay showed that introduction of AS1-S increased the number of mRNA that co-immunoprecipitated with bovine hnRNPM in MDBK cells. These results suggested that AS1-S could alter the interaction between hnRNPM and host mRNAs, potentially interfering with cellular functions during the initial phase of mRNA maturation in the nucleus. Since most of the identified mRNAs that exhibited increased binding to hnRNPM were correlated with the KEGG term “Pathways in cancer”, AS1-S may affect proliferation and expansion of BLV-infected cells and contribute to tumor progression.

**Importance:** BLV infects bovine B cells and causes malignant lymphoma, resulting in severe economic losses in the livestock industry. Due to its low incidence rate and long latent period, the molecular mechanisms underlying the progression to lymphoma remain enigmatic. Several non-coding RNAs, such as miRNA and lncRNA, have recently been discovered in the BLV genome and the relationship between BLV pathogenesis and these non-coding RNAs is attracting attention. However, most of the molecular functions of these transcripts remain un-identified. To the best of our knowledge, this is the first report describing a molecular function for the BLV-derived lncRNA AS1-S. The findings reported herein reveal a novel mechanism underlying BLV pathogenesis that could provide important insights for not only BLV research but also comparative studies of retroviruses.

## Introduction

Viruses utilize several strategies to maintain latent infection while evading host immune responses (1). One of the strategies is the expression of functional non-coding RNA (ncRNA); the viral genome encodes ncRNA to modulate gene expression of host or viral genes in infected cells (2–5). Since ncRNAs can evade host immune responses due to their lack of antigenicity, they are an effective tool for maintaining latent infection. A representative example of ncRNA is miRNA, which is a class of ∼21 nucleotide small RNAs that repress gene expression by increasing RNA degradation or inhibiting translation and alters numerous cellular processes in infected cells (6). Additionally, ncRNAs include long non-coding RNA (lncRNA), a class of RNA with lengths of > 200 nucleotides that do not encode protein; most lncRNAs regulate various cellular functions by interacting with the genome and RNA binding proteins (3, 4, 7).

Bovine leukemia virus (BLV), which belongs to the genus Deltaretrovirus of the family Retroviridae and causes a malignant B cell lymphoma in cattle, encodes several ncRNAs including miRNA and lncRNA (8–11). BLV expresses structural and accessary proteins and aggressively expands during the early stage of infection. However, upon establishing a latent state during chronic infection in BLV-infected cells, BLV expresses few viral antigens encoded in the sense strand of the genome (12, 13). In contrast to the protein-coding transcripts, expression of miRNAs and antisense transcripts, all of which are lncRNA, are continuously active during latent infection (9, 11, 14). Among these ncRNAs, the miRNAs are likely to be responsible for BLV pathogenesis (15, 16). The BLV-derived lncRNA AS1 consists of two isoforms, AS1-L and AS1-S, and is highly expressed in the nucleus of BLV-infected cells (11, 14). Although the function of the AS1-S remains unknown, it is hypothesized to play important roles in the BLV lifecycle because of its continuous expression.

Since most lncRNAs function by interacting with RNA-binding proteins, AS1-S potentially interacts with several proteins in infected cells. Interactions between AS1-S and its binding partners might play pivotal roles in modulating the cellular environment during latent infection or in the progression of lymphoma. Therefore, identifying the binding partners of AS1-S and clarifying its function is highly significant for BLV research. Furthermore, the obtained findings are expected to be widely applicable to understanding the survival strategies of retroviruses. In this study, we established a bovine cell line stably expressing AS1-S and identified bovine heterogeneous nuclear ribonucleoprotein M (hnRNPM), an RNA-binding protein located in the nucleus, as the binding partner for AS1-S. The AS1-S-expressing cells altered the variety of mRNAs that co-immunoprecipitated with bovine hnRNPM, implying that AS1-S modifies cellular function by altering interactions between hnRNPM and host mRNAs in the nucleus.

## Results

### Construction and evaluation of AS1-S RNA expressing MDBK cells

To evaluate functional changes in cells associated with AS1-S, we established MDBK cells expressing AS1-S under control of the CAG promoter or its internal 3’LTR promoter (Fig. S1A and B in the supplemental material). Localization of AS1-S in these cells were confirmed using real-time RT-PCR, which indicates that the localization of AS1-S in transgenic MDBK cells differed from that in the BLV-infected B cell line BL3.1. Specifically, AS1-S in BL3.1 cells is mainly located in the nucleus (approximately 70%), while transgenic AS1-S in MDBK cells is mainly located in the cytoplasm (the ratio of nuclear RNA was < 30%) (Fig. S1C in the supplemental material). Notably, the ratio of nuclear AS1-S RNA in MDBK CAG AS1-S cells was quite low (< 10%) relative to MDBK 3’LTR AS1-S cells (approximately 30%), which is consistent with a previous report indicating that an antisense RNA driven by a strong promoter was located in the cytoplasm (19). Localization of nuclear (U6) and cytoplasmic (TYR) RNA controls in the transfected cells were consistent with BL3.1 cells, supporting the validity of the experiment (Fig. S1C in the supplemental material). These results indicated that MDBK 3’LTR AS1-S cells were more suitable than MDBK CAG AS1-S cells for subsequent experiments, although the ratio of nuclear AS1-S RNA in MDBK 3’LTR AS1-S cells was lower than in BL3.1 cells.

### Transcriptome analysis of transfected MDBK cells

The transfected MDBK cells were subjected to transcriptome analysis in an effort to identifying differentially expressed genes (DEGs). A PCA plot showed that MDBK 3’LTR AS1-S cells were clustered separately from the mock and parental cells. However, MDBK mock cells also exhibited an altered expression profile compared to the parental MDBK cells due to insertion of the empty vector or G418 selection (Fig. 1A). Subsequently, the mRNA expression profile of MDBK 3’LTR AS1-S cells (n=2) was compared with MDBK mock and parental cells (n=4) to identify AS1-S specific effects. The resultant differential expression analysis identified 83 DEGs in MDBK 3’LTR AS1-S cells, with 28 and 55 up- and down-regulated genes, respectively (Fig. 1B–D, DEGs are listed in Table S2 in the supplemental material). The identified DEGs were then subjected to GO analysis and the result showed that molecular function (MF) terms including “signaling receptor binding”, “receptor ligand activity”, “signaling receptor activator activity”, and “signaling receptor regulator activity” were significantly enriched (Fig. 1E, results of the GO analysis are listed in Table S3 in the supplemental material). Moreover, the biological process (BP) terms “nervous system development”, “neurogenesis”, “cell migration”, “cell motility”, and “localization of cell” were enriched, although the p-values were not significant (p > 0.05) (Fig. 1F). In this experiment, no significant result was obtained from KEGG pathway enrichment analysis (data not shown).

**Fig. 1.**
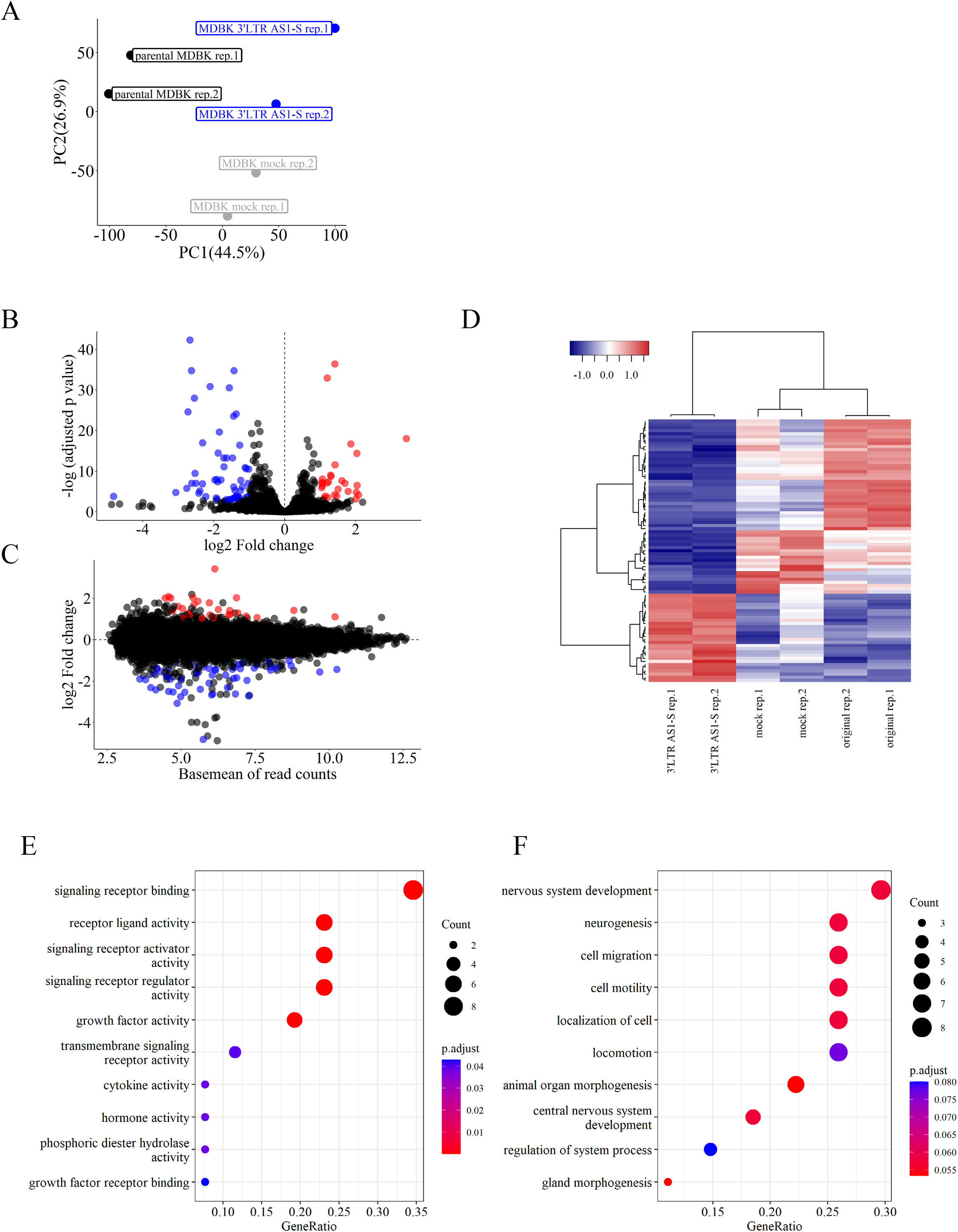
(A) Principal component analysis (PCA) plot of the transcriptome analysis using MDBK 3’LTR AS1-S, MDBK mock, and parental MDBK cells. The X and Y axes show principal components 1 and 2, and percentages in parentheses indicate their respective contributions to the overall variability. (B–D) Volcano plot (B), smear plot (C) and heatmap (D) of the transcriptome analysis results. Significant differentially expressed genes (DEGs) identified between MDBK 3’LTR AS1-S (n=2) and MDBK mock and parental cells (n=4) are shown as red and blue (up- and down-regulated in MDBK 3’LTR AS1-S cells, respectively). Significant DEGs were defined based on the following criteria: |fold change| ≥ 2, exactTest raw p-value < 0.05. (E, F) Gene ontology (GO) analysis of significant DEGs. The top ten enriched terms in molecular function (MF) (F) and biological process (BP) (G) are shown. The size of the bubbles indicates the “gene count”, and the color of the bubbles indicates the “adjusted p-value”. Adjusted p-value < 0.05 is defined as significant.

### Identification of a host-derived protein that binds to the AS1-S RNA probe

Since introduction of AS1-S alters gene expression in MDBK cells, we aimed to identify a binding partner of AS1-S using an AS1-S RNA probe and a BL3.1 cell lysate. An RNA-protein pull-down assay resulted in the observation of three bands that were specific to the sample obtained from the AS1-S RNA probe; the size of the bands were > 100 kDa, ∼ 70 kDa, and < 20 kDa (Fig. 2A). The three bands were analyzed by LC-MS and the ∼ 70 kDa band was predicted to be bovine hnRNPM. To confirm the result obtained with LC-MS, western blotting analysis using an anti-hnRNPM monoclonal antibody was performed, and the result showed that a bovine hnRNPM band was specifically observed at ∼ 70 kDa in the sample from the AS1-S RNA probe (Fig. 2A). These results indicated that AS1-S RNA physically interacts with bovine hnRNPM.

**Fig. 2.**
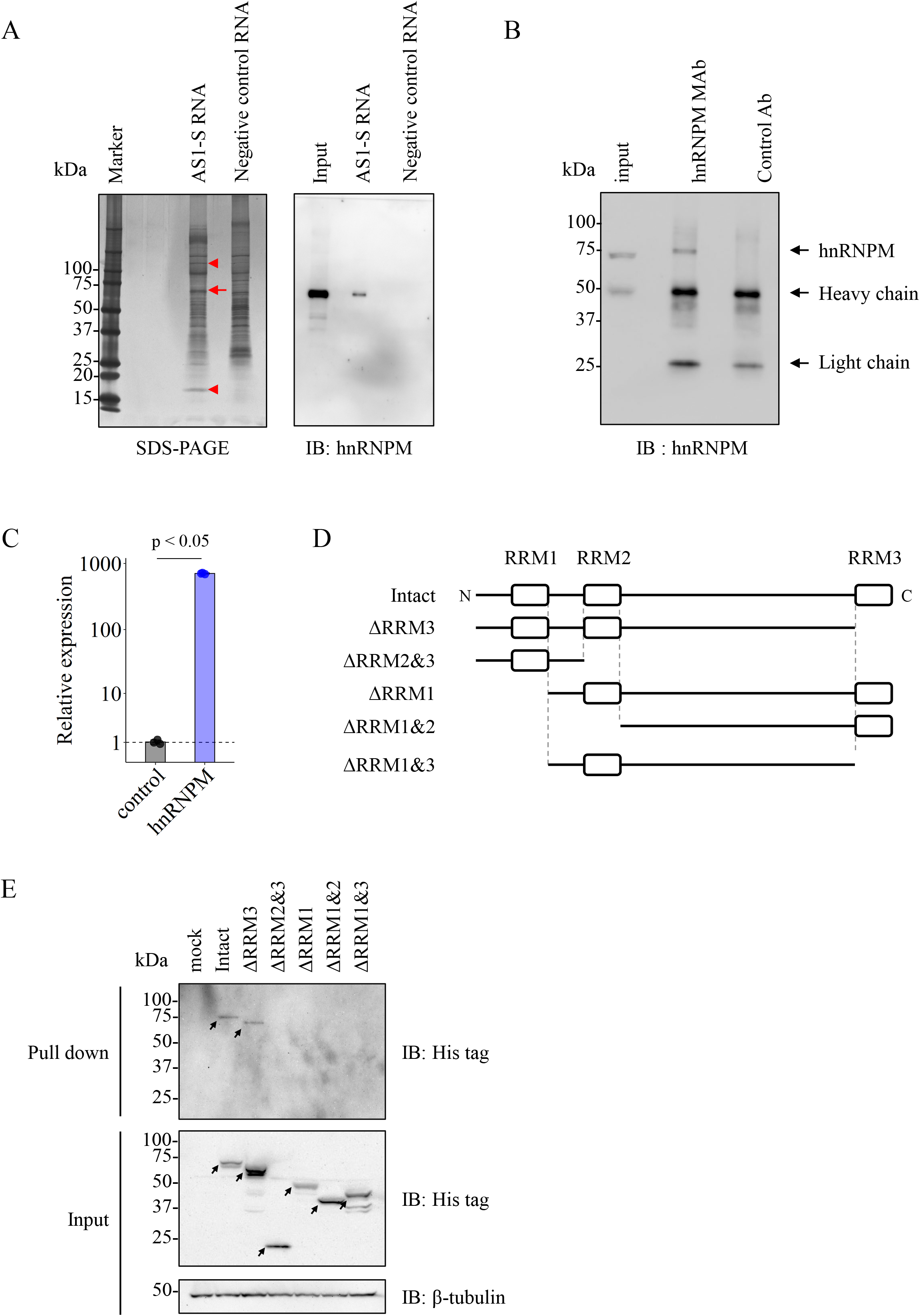
(A) Pull-down assay results using RNA probes. Biotinylated AS1-S or control RNA probes were mixed with a BL3.1 cell lysate and pulled-down samples were subjected to SDS-PAGE (silver staining) (left), followed by western blotting (right). The red arrow shows the band that is predicted as bovine hnRNPM using LC-MS. Two arrow heads indicate bands that are specific for the AS1-S RNA probe but did not produce a significant result with LC-MS. (B) Western blotting analysis of RIP samples. RNA-protein complexes were immunoprecipitated from BL3.1 cell lysates using an anti-hnRNPM or a control monoclonal antibody, and then subjected to western blotting. Heavy and light chains observed in the immunoprecipitated samples were derived from the antibodies used for the immunoprecipitation procedure. (C) Quantification of AS1-S RNA in RNA immunoprecipitated samples. RNA was purified from an RNA-protein complex obtained by RNA immunoprecipitation, followed by real-time RT-PCR. The result is shown as expression levels relative to the control sample. Data are presented as the mean + standard deviation (n = 3), and a t-test was performed for statistical analysis. p < 0.05 was defined as statistically significant. (D) Schematic diagrams of bovine hnRNPM protein and the constructed deletion mutants. White boxes indicate RNA recognition motifs (RRM). All constructs were fused with a his-tag sequence and inserted into the expression plasmid pCAG neo. (E) Pull-down assay results using AS1-S RNA probe and recombinant hnRNPMs. Recombinant proteins were expressed in 293T cells and the obtained cell lysates were mixed with a biotinylated AS1-S RNA probe, and subsequently assessed by western blotting. Arrows indicate the predicted sizes of the recombinant proteins.

### Confirmation of the interaction between AS1-S and hnRNPM in bovine B cells

To confirm whether the interaction between AS1-S and bovine hnRNPM occurs in BLV-infected B cells, an hnRNPM-RNA complex was extracted from BL3.1 cells using an RNA immunoprecipitation (RIP) assay. The RIP assay result indicated that the anti-hnRNPM antibody specifically immunoprecipitated bovine hnRNPM (Fig. 2B), and real-time RT-PCR showed that the amount of the AS1-S RNA immunoprecipitated with anti-hnRNPM was approximately 700 times greater than with the control antibody (Fig. 2C). These results suggested that the AS1-S RNA interacts with hnRNPM in BLV-infected B cells.

### Identification of bovine hnRNPM regions responsible for binding AS1-S

Bovine hnRNPM contains three RNA recognition motifs (RRM) in its amino acid sequence. To identify the regions responsible for binding AS1-S, bovine hnRNPM deletion mutants were constructed and subjected to the pull-down assay (mutant constructs are shown in Fig. 2D). The pull-down assay using the mutant constructs and the AS1-S RNA probe showed that only the full-length and ΔRRM3 hnRNPMs, both of which include RRM1 and RRM2, bound the AS1-S RNA probe (Fig. 2E). This result indicated that both RRM1 and RRM2 are required for the interaction between bovine hnRNPM and AS1-S.

### Knock down of hnRNPM expression in BL3.1 cells

To confirm whether knock down of hnRNPM affects the expression of viral proteins in BL3.1 cells, siRNAs targeting bovine hnRNPM were introduced into BL3.1 cells. Western blotting analysis showed that the expression of hnRNPM was reduced by the introduction of two siRNAs targeting hnRNPM (sihnRNPM #1 and #2). However, the expression of viral proteins such as gp51 and p24 was not affected in these cells (Fig. 3).

**Fig. 3.**
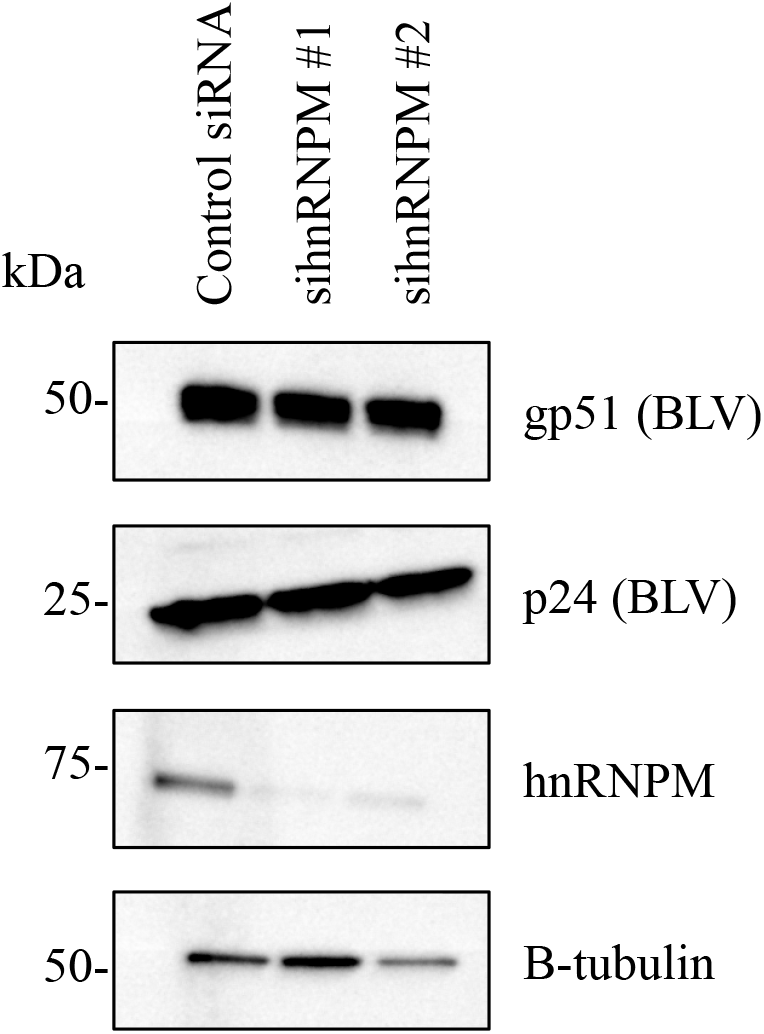
Western blotting analysis of BL3.1 cells transfected with siRNAs. Two siRNAs against bovine hnRNPM and a scrambled control were introduced into BL3.1 cells and the expression of each protein was visualized using monoclonal antibodies.

### Analysis of the alternative splicing events

Because hnRNPM is involved in alternative splicing (23, 24), the RNA-seq data used for the transcriptome analysis in Fig. 1 were subjected to alternative splicing analysis to determine whether the introduction of AS1-S affects mRNA splicing. The comparison was performed under the same conditions as the transcriptome analysis (MDBK 3’LTR AS1-S (n=2) vs MDBK mock and parental cells (n=4)), and 39 of 5758 events were identified as significantly different between the two groups; the number of the significant SE, A5SS, A3SS, MXE, and RI events were 15 of 3873, 3 of 414, 13 of 959, 4 of 259, and 4 of 253, respectively (Fig. S2, rMATS output are listed in Table S4 in the supplemental material). However, no GO terms were enriched among these 39 genes (data not shown).

### Comprehensive analysis of hnRNPM-binding RNAs in transfected cells

Because AS1-S interacted with the RRMs in hnRNPM, it is possible that the interaction between AS1-S and hnRNPM affects interactions between hnRNPM and endogenous RNAs. To evaluate whether AS1-S interfered with interactions between hnRNPM and host-derived RNA, RNAs interacting with hnRNPM were comprehensively analyzed by RIP-seq. The RIP assay using transfected cells showed that the amount of AS1-S RNA immunoprecipitated with the anti-hnRNPM antibody was approximately 6 times greater than with the control antibody in MDBK 3’LTR AS1-S cells (Fig. 4A), indicating that exogenous AS1-S bound to hnRNPM in MDBK cells. The samples were subsequently subjected to RNA-seq analysis (RIP-seq), followed by visualization as read counts for each mRNA (Fig. 4B). The results showed that the number of mRNAs that co-immunoprecipitated with hnRNPM (read counts > 10, hnRNPM/control ratio > 2.0) in MDBK 3’LTR AS1-S and MDBK mock cells were 5602 and 1652, respectively (shown as red dots in Fig. 4B, the RIP-seq results are listed in Table S5 and S6 in the supplemental material). Evaluation of the gene lists indicated that 995 genes were observed in both MDBK 3’LTR AS1-S and MDBK mock cells, and 4607 of 5602 genes (82.2%) were observed only in MDBK 3’LTR AS1-S cells (Fig. 4C). This result suggests that the introduction of AS1-S increased the variety of mRNAs that co-immunoprecipitated with hnRNPM.

**Fig. 4.**
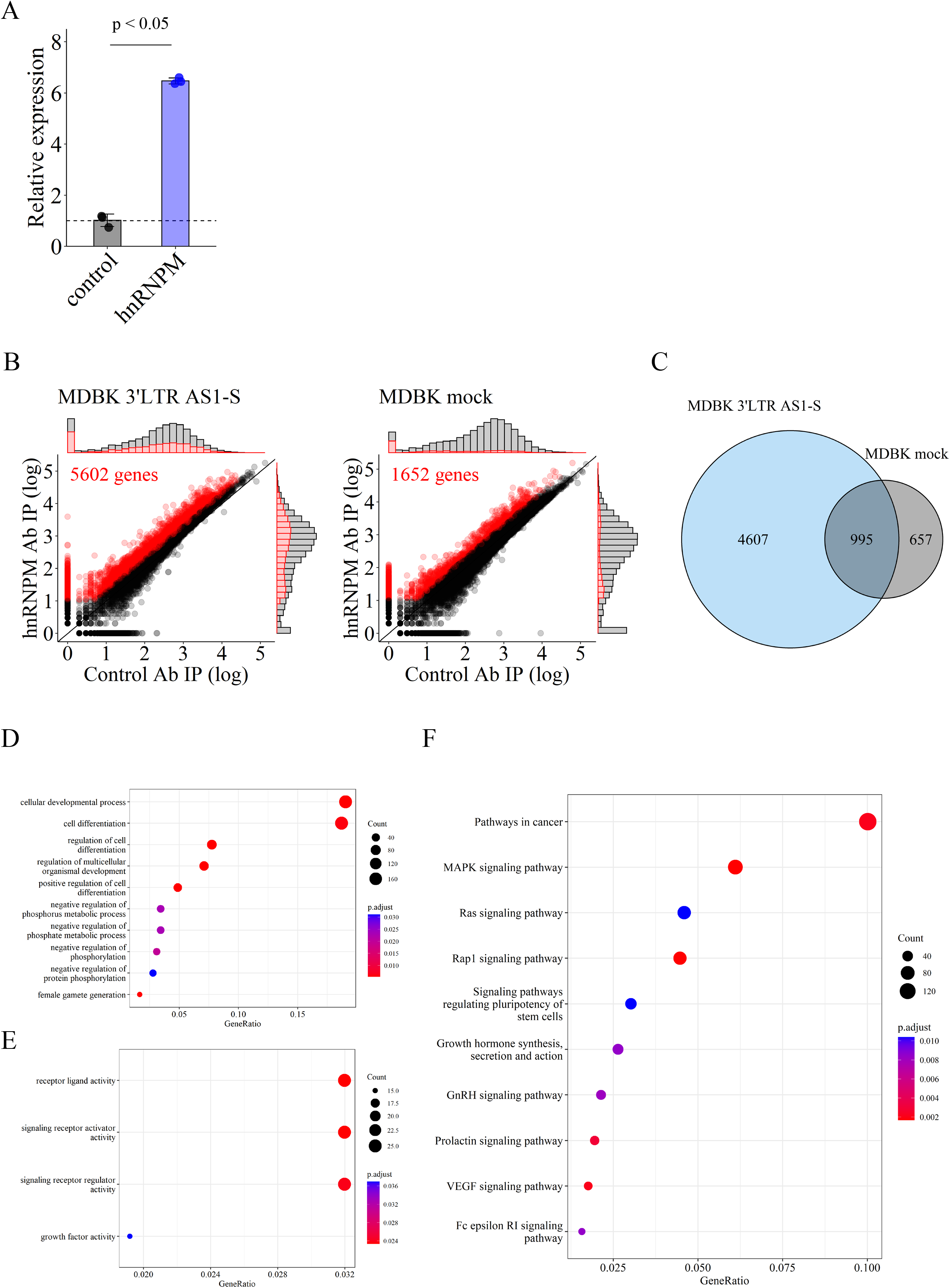
(A) Quantification of AS1-S RNA in RNA immunoprecipitated samples from MDBK 3’LTR AS1-S cells. RNA was purified from an RNA-protein complex obtained by RNA immunoprecipitation, followed by real-time RT-PCR. The result is shown as expression levels relative to the control sample. Data are presented as the mean + standard deviation (n=3), and a t-test was performed for statistical analysis. p < 0.05 was defined as statistically significant. (B) Scatter plots showing sequencing reads obtained by RNA immunoprecipitation. The X and Y axes show read counts obtained by RIP-seq from control MAb and anti-hnRNPM MAb, respectively. Genes that meet the following criteria were defined as hnRNPM-binding RNAs and are shown as red dots: read counts > 10, hnRNPM/control ratio > 2.0. (C) Venn diagram showing the overlapping hnRNPM-binding genes in MDBK 3’LTR AS1-S and MDBK mock cells. (D–F) Gene ontology (GO) analysis of the 4607 hnRNPM-binding genes only detected in MDBK 3’LTR AS1-S cells. Molecular function (MF) (D), biological process (BP) (E) and KEGG pathway enrichment analysis (F) are shown separately. The size of the bubbles indicates the “gene count”, and the color of the bubbles indicates the “adjusted p-value”. Adjusted p-value < 0.05 is defined as significant.

To elucidate common functions of the 4607 mRNAs identified in Fig. 4C, the gene list was subjected to GO analysis. The result indicated that the BP terms “cellular developmental process” and “cell differentiation”, as well as the MF terms “receptor ligand activity”, “signaling receptor activator activity”, and “signaling receptor regulator activity” were significantly enriched (Fig. 4D and E, results of enrichment analysis are listed in Table S7 in the supplemental material). Furthermore, KEGG pathway enrichment analysis showed that most of the identified genes were related to pathways for cell proliferation, such as “pathways in cancer”, “MAPK signaling pathway”, “Ras signaling pathway”, and “Rap1 signaling pathway” (Fig. 4F and S3, results of the enrichment analysis are listed in Table S8 in the supplemental material).

### Analysis of mRNA distribution in AS1-S-expressing cells

Because several heterogeneous nuclear ribonucleoproteins are involved in the nuclear export of mRNAs (25, 26), we measured nuclear and cytoplasmic RNA in MDBK 3’LTR AS1-S and MDBK mock cells and comprehensively evaluated the distribution of RNA to confirm whether the interaction between AS1-S and hnRNPM affected mRNA translocation. Read counts of nuclear and cytoplasmic RNAs showed that the nuclear/cytoplasmic RNA ratio in AS1-S-transfected and mock cells was similar (Fig. S4A, the count matrix is listed in Table S9 in the supplemental material). Moreover, the 4607 mRNAs identified in Fig. 4C exhibited a similar nuclear/cytoplasmic RNA ratio in AS1-S-transfected and mock cells (Fig. S4B in the supplemental material). These results indicated that introduction of AS1-S did not affect the nuclear export of mRNAs.

### Analysis of mRNA distribution and hnRNPM-binding RNAs in BL3.1 cells

We also analyzed mRNA distribution and hnRNPM-binding RNAs in BL3.1 cells as supplemental data although no proper control, such as BLV-negative B cells, could not be prepared. Scatter plots of nuclear and cytoplasmic RNA in BL3.1 were largely identical to that of MDBK 3’LTR AS1-S and mock cells (Fig. S5A, the count matrix is listed in Table S10 in the supplemental material). In contrast, the RIP-seq results differed remarkably from that of MDBK-derived cells; the number and amount of RNAs that co-immunoprecipitated with the anti-hnRNPM was greater than with the control antibody in BL3.1 cells (Fig. S5B, the RIP-seq results are listed in Table S11 in the supplemental material). Western blotting analysis showed that the expression of hnRNPM was almost equal among the same number of BL3.1 and MDBK-derived cells (Fig. S5C in the supplemental material), although the expression of beta-tubulin in BL3.1 cells was less than in MDBK cells. This result indicates that the difference observed in RIP-seq might be attributable to the cell type or BLV infection.

## Discussion

In this study, we identified bovine hnRNPM as the binding partner of the BLV-derived lncRNA, AS1-S. Pull-down assays using recombinant hnRNPMs showed that both RRM1 and 2 were responsible for binding AS1-S (Fig. 2). In addition, we found that the introduction of AS1-S increased the variety of RNA that co-immunoprecipitated with bovine hnRNPM in MDBK cells. Moreover, the number of mRNAs that formed a complex with hnRNPM in AS1-S-transfected cells was approximately three times greater than in mock cells, and KEGG pathway enrichment analysis suggested that most of the identified RNAs were correlated with the KEGG term “Pathways in cancer” (Fig. 4). These results implied that AS1-S could alter interactions between hnRNPM and mRNAs and potentially alter the proliferation and expansion of infected cells, which could be novel mechanisms for the progression of lymphoma. To our best knowledge, this is the first report attributing molecular functions to BLV lncRNA.

Since AS1-S bound to the RRM1 and 2 of hnRNPM, we hypothesized that AS1-S might physically interfere with interactions between hnRNPM and host RNAs. Contrary to our expectation, introduction of AS1-S increased the number of mRNAs co-immunoprecipitating with hnRNPM in MDBK cells (Fig. 4B and C). As a consequence, AS1-S was hypothesized to alter interactions between hnRNPM and host mRNAs that result in the regulation of the expression of several genes during the initial transcription processes occurring in the nucleus. Although it remains unknown how AS1-S RNA increases interaction between hnRNPM and several mRNAs and whether the identified mRNAs directly bind to hnRNPM, liquid-liquid phase separation (LLPS) could be implemented to explain the mechanism. Previously, Unfried and Ulitsky proposed that lncRNAs can form biomolecular condensates in cells and that biomolecular condensates can facilitate enzyme activities by locally increasing the concentration of enzymes and substrates (27). Their review suggested that such reversible condensates compensate for the low expression levels of lncRNAs (27). Therefore, AS1-S RNA could function in the same manner to form a condensate with hnRNPM, thereby altering hnRNPM-RNA interaction. Moreover, the present RNA-protein pull-down data suggests that the AS1-S RNA probe could bind hnRNPM as well as other host-derived proteins (Fig. 2A). Therefore, it is possible that several unknown molecules also support the formation of biomolecular condensates and increase interactions between hnRNPM and mRNAs. This hypothesis is consistent with a previous report that demonstrated that AS1-S RNA was observed as small nuclear dots by *in situ* hybridization (14).

In the present study, we could not identify specific genes or pathways that are directly affected by the interaction between AS1-S and hnRNPM since numerous factors were identified by RIP-seq and KEGG pathway enrichment analysis (Fig. 4 and S3 in the supplemental material). However, these analyses showed that the KEGG pathway “Pathways in cancer” was significantly enriched (Fig. 4F), increasing the possibility that AS1-S acts as a determinant for the abnormal proliferation of BLV-infected cells. Generally, pathways correlating with cancer primarily facilitate cell proliferation, implying that AS1-S could contribute to the prolonged lifespan of infected-cells, thereby increasing the probability of acquiring lethal mutations leading to the progression of lymphoma. This finding could be helpful in unveiling novel molecular mechanisms underlying tumorigenesis in BLV-infected cells.

Several factors have been reported to be important determinants related to the pathogenesis of BLV, including viral factors with transcriptional activity, defined by interactions between long terminal repeat (LTR) and the TAX protein (28–30). Since transcriptional activity strongly affects BLV replication, it is expected to contribute to the expansion of BLV during the initial infection step. On the other hand, since latently infected BLV expresses no viral proteins, the functions of LTR and TAX cannot completely explain the behavior of the virus during late stage infection. Moreover, host-derived factors such as BoLA-DRB3 polymorphisms (31, 32) and immune exhaustion during BLV infection (33, 34) were well characterized in an attempt to elucidate this concern. In addition to these reports, Durkin et al. provided the novel insight that antisense transcripts from the BLV provirus genome are stably expressed during latent infection and may play pivotal roles in the BLV life cycle (11). Rosewick et al. subsequently showed that BLV antisense transcripts produced chimeric transcripts with host mRNA, a critical step in the progression of lymphoma (35). Herein, we clarified the binding partner of the AS1-S RNA and demonstrated that it functions as a molecular modulator of bovine hnRNPM RNA-interaction. This is the first report describing the molecular function of the antisense transcript encoded by BLV, an insight that is expected to contribute to future research into BLV pathogenesis.

Interestingly, human and mouse hnRNPM contribute to the innate immune response against pathogens (36, 37). Additionally, other reports showed that alternative splicing events modulated by hnRNPM correlate with tumor progression (38, 39). Regarding the former, Cao et al. reported that hnRNPM was translocated from the nucleus to cytoplasm in response to infection with RNA viruses, resulting in suppression of innate immune responses by antagonizing RNA sensors (36). West et al. also reported that hnRNPM modulates splicing of IL6 mRNA and controls innate immune responses against bacteria (37). In our study, however, no GO terms relating to immune responses were significantly enriched in both RIP-seq and transcriptome analysis (Fig. 1 and 4), and the splicing pattern of IL6 was not found to be altered in the rMAST analysis (Fig. S2 and Table S4 in the supplemental material). Therefore, it remains unknown whether the interaction between AS1-S and bovine hnRNPM affects the immune response in BLV-infected cells. Hypothetically, since BLV expresses no viral antigens during the latent phase of infection, suppression of immune responses might not be required in latently infected cells, and the interaction of AS1-S and hnRNPM could have another role other than immune suppression.

One of the limitations of this study is that we were not able to perform gain of function and loss of function analyses using bovine B cells. Currently, bovine B cell lines that are free from BLV infection are not available; therefore, it was impossible to perform a gain of function analysis by introducing AS1-S into bovine B cells. Since all the experiments in this study used kidney derived cells, differences related to cell type could introduce some biases; for example, the AS1-S RNA expressed in MDBK cells was mainly located in the cytoplasm despite its expression being driven by the 3’LTR promoter (Fig. S1C in the supplemental material). In addition to the concerns pertaining to cell origin, the low transfection efficiency in bovine cell lines restricts implementation of certain experiments (40, 41). Since a transient expression strategy was not suitable for our experiments, G418 selection was used to prepare stably expressing cells, a procedure that has the potential to introduce additional biases into the results. In fact, the PCA plot showed that both the MDBK 3’LTR AS1-S and MDBK mock cells exhibited altered transcriptome profiles relative to the parental MDBK cells (Fig. 1A). Moreover, the transcriptome analysis of MDBK cells expressing AS1-S identified significant DEGs, but the obtained GO terms were not directly related to BLV pathogenesis (Fig. 1B–F). Thus, these phenotypic changes can be caused by the biases described above. In addition, it should also be noted that we attempted to knock down the expression of AS1-S RNA in BL3.1 cells; however, siRNAs and antisense oligos were unable to successfully downregulate the expression of AS1-S RNA, although we successfully knocked down hnRNPM expression using siRNA (Fig. 3). The low transfection efficiency of BL3.1 cells may be insufficient to knock down the expression of AS1-S RNA located in the nucleus. Taken together, limitations related to bovine cells are an obstacle to BLV research. Thus, the establishment of novel bovine B cell lines free from BLV infection and the development of new transfection methodologies that are capable of efficiently transfecting bovine cells is required to advance BLV research.

There are some outstanding questions that remain to be clarified after this study. For example, although LC-MC successfully identified bovine hnRNPM as the binding partner for AS1-S, the other bands (> 100 kDa and < 20 kDa) could not be identified (Fig. 2A). Thus, there are additional protein factors involved in mediating AS1-S functions, and it is possible that these factors function independently or cooperatively with hnRNPM. Furthermore, since our study mainly focused on the protein-RNA interaction between AS1-S and hnRNPM, we could not rule out the possibility that the interaction of these molecules inhibits protein-protein interactions involving hnRNPM. Additionally, although we identified that RRM1 and 2 of hnRNPM were regions required for binding AS1-S RNA, the biological importance of each RRM has not been clarified. Thus, it remains to be elucidated how the RNA-binding properties of these RRMs affects the biological functions of bovine hnRNPM. Since the functions of bovine hnRNPM have not been elucidated at all, future research aiming to clarify the molecular functions of bovine hnRNPM could also provide new insights for BLV research.

It is worth mentioning that the RIP-seq result of BL3.1 cells was drastically different from that of MDBK-derived cells (Fig. S5 in the supplemental material). Western blotting analysis showed that the expression levels of bovine hnRNPM were similar in BL3.1 and MDBK-derived cells; however, the RIP-seq data showed that the number of mRNAs co-immunoprecipitating with hnRNPM in BL3.1 cells was much greater than in MDBK cells (Fig. 4 and S5 in the supplemental material). One hypothesis for this result is that the RNA binding ability of hnRNPM is more functionally important in B cells and strong, and therefore the variety of hnRNPM-interacting RNAs in B cells is more diverse than in other cells. This hypothesis may explain why only B cells undergo tumorigenesis following BLV infection. The other hypothesis is that persistent BLV infection of BL3.1 cells alters the function of bovine hnRNPM more drastically than exogenous AS1-S RNA. Since knock down analysis showed that siRNAs against hnRNPM did not alter the expression of viral proteins (gp51 and p24, Fig. 3) in BL3.1 cells, the interaction between AS1-S and hnRNPM might not be important for the transcription of viral sense RNA, but rather the maintenance of B cell infection. To confirm the hypothesis, the development of a novel bovine B cell line is necessary.

Herein, we showed that the retroviral lncRNA AS1-S bound to the host hnRNPM and modulated its RNA-binding profile. This novel insight is expected to bring a new perspective to research into retroviral antisense transcripts. Regarding retroviral antisense transcripts, the HBZ gene encoded in human T-cell leukemia virus (HTLV), which also belongs to the genus Deltaretrovirus of the family Retroviridae, is well characterized (42). While HBZ functions as a protein, its RNA form has bimodal functions in HTLV-infected cells (5); HBZ protein suppresses TAX-mediated transcription through the 5’LTR, whereas HBZ RNA promotes cell proliferation and inhibits apoptosis (43). The function of HBZ RNA was reported to involve the upregulation of the expression of many genes related to the cell cycle, proliferation and survival by interacting with their promoter sequences (44). Additionally, it was reported that HBZ RNA directly affects interactions between RNA polymerase and the viral LTR promoter by displacing a transcriptional factor (45). Thus, diverse roles of HBZ RNA have been reported, and our finding that the retroviral lncRNA, AS1-S, interacts with host hnRNPM has broadened the understanding of viral genome function in retrovirus lifecycle. Evaluation of the similarities and differences between HBZ and AS1-S might reveal important insights relevant to BLV research as well as comparative studies of retroviruses.

## Materials and methods

### Cells

Madin-Darby bovine kidney (MDBK) cells were maintained in Eagle’s medium (Nissui Pharmaceutical, Tokyo, Japan) supplemented with 5% heat-inactivated fetal bovine serum (FBS) (Thermo Fisher Scientific, Waltham, MA, USA), 100 units/mL penicillin, 100 μg/mL streptomycin (Sigma-Aldrich, St Louis, MO, USA), and 2 mM L-glutamine (Nacalai Tesque, Kyoto, Japan). 293T cells (ATCC CRL-3216) were maintained in Dulbecco’s modified Eagle’s medium (DMEM) (Nissui Pharmaceutical) supplemented with 10% FBS along with 100 units/mL penicillin and 100 μg/mL streptomycin. BL3.1 cells (ATCC CRL2306), which are persistently infected with BLV (previously characterized (17, 18)), were maintained in RPMI1640 GlutaMAX (Thermo Fisher Scientific) supplemented with 10% FBS along with 100 units/mL penicillin and 100 μg/mL streptomycin.

### Construction of plasmids

An expression plasmid encoding BLV AS1-S under control of the CAG promoter was previously reported (pCAG AS1-S: Nishimori et al., in press); briefly, 571 bp of AS1-S cDNA derived from FLK-BLV (LC164083.1) was inserted downstream of the CAG promoter (Fig. S1A). Because promoter sequences alter the localization of antisense RNA encoded in deltaretrovirus (19), an expression plasmid without the CAG promoter was also constructed; the CAG promoter sequence was removed from pCAG AS1-S by digesting with *Nde*I and *Sal*I, and the resultant fragment lacking the CAG promoter was treated with Blunting high (TOYOBO, Osaka, Japan), followed by self-ligation. The constructed plasmid was designated as p3’LTR AS1-S because the AS1-S RNA is transcribed under control of its internal 3’LTR promoters IRF and DAS (11).

To construct a plasmid for synthesizing an AS1-S RNA probe, the AS1-S sequence was obtained from pCAG AS1-S by digesting with *Hin*dIII and *Pst*I. The fragment was subsequently inserted into the pSPT19 plasmid (designated as pSPT19 AS1-S) using the same restriction enzymes.

To construct expression plasmids encoding recombinant hnRNPMs, a complete cDNA sequence of bovine hnRNPM (NM_001191223) was obtained from genomic DNA of MDBK cells using PrimeScript RT reagent Kit (TaKaRa, Shiga, Japan), followed by a PCR reaction using PrimeSTAR Max DNA Polymerase (TaKaRa) with primers no. 1 and 4 (all primers are listed in Table S1 in the supplemental material). The PCR conditions were as follow; 35 cycles of 98 °C for 10 s, 55 °C for 5 s, and 72 °C for 2 min. The amplicon was then cloned into the *Eco*RI and *Not*I recognition site of the expression plasmid pCAG neo using HD Cloning Kit (TaKaRa). To construct deletion mutants of bovine hnRNPM, sequence fragments were amplified from the complete hnRNPM cDNA using PrimeSTAR Max DNA Polymerase (TaKaRa) with primer sets no. 1–6; the PCR conditions were same as those used for amplification of the complete hnRNPM sequence. The resultant fragments were cloned into the *Eco*RI and *Not*I recognition sites of pCAG neo using HD Cloning Kit (TaKaRa). All recombinant proteins were fused with a his-tag and expression was confirmed using an anti-his tag antibody.

### Design of small interfering RNA (siRNA)

Two pairs of stealth RNAi targeting bovine hnRNPM were designed and purchased from Thermo Fisher Scientific. The paired sequences are described below; sihnRNPM1: 5’-GGCAGUCACUUAAAGACCUGGUUAA-3’ (sense) and 5’-UUAACCAGGUCUUUAAGUGACUGCC-3’ (antisense), and sihnRNPM2: 5’-GAGGUAACAUACGUGGAGCUCUUAA-3’ (sense) and 5’-UUAAGAGCUCCACGUAUGUUACCUC-3’ (antisense). The scrambled sequences 5’-GGCCUCAUUAAGACAUCGGUGAUAA-3’ (sense) and 5’-UUAUCACCGAUGUCUUAAUGAGGCC-3’ (antisense) were used as a control.

### Transfection or nucleofection and establishment of stable cell lines

Constructed plasmids encoding AS1-S were introduced into MDBK cells using Amaxa cell line Nucleofector kit R (Lonza, Kanagawa, Japan) with Amaxa Nucleofector II system in accordance with the manufacturer’s instructions. Briefly, 1 µg of plasmid DNA and 1 × 10^6^ cells were mixed with the nucleofector reagent, followed by nucleofection using the X-001 program. After nucleofection, the cells were subjected to G418 selection.

293T cells were transfected with the plasmids encoding recombinant bovine hnRNPMs using polyethylenimine (PEI). Briefly, 1 µg of each plasmid was mixed with OPTI-MEM (Thermo Fisher Scientific) containing 10 µL of PEI reagent (2 mg/mL concentration of PEI MAX MW 40,000: Polysciences, Warrington, PA, USA) and then transfected into 293T cells grown to confluency in 6-well plates. Transfected cells were harvested at 72–96 hr post-transfection and subjected to subsequent experimentation.

siRNAs targeting bovine hnRNPM were introduced into BL3.1 cells using Amaxa cell line Nucleofector kit V (Lonza) in accordance with the manufacturer’s instructions. Briefly, 10 µL of 20 µM siRNA and 1 × 10^6^ cells were mixed with the suspension buffer and subsequently subjected to the nucleofection using the O-017 program. At 72 hr post-nucleofection, cells were harvested, washed with PBS and then subjected to western blotting analysis.

Establishment of stable cell lines was performed in accordance with a previous report (Nishimori et al., in press); briefly, expression plasmids encoding AS1-S (pCAG AS1-S and p3’LTR AS1-S) were introduced into MDBK cells, and the transfected cells were maintained in the presence of 1,000 μg/mL of G418 (Thermo Fisher Scientific). As a control, a stable cell line introduced with an empty vector (pCAG neo) was established in the same manner. The established cells were designated as MDBK CAG AS1-S, MDBK 3’LTR AS1-S, and MDBK mock.

### RNA-protein pull-down assay and LC-MS

An RNA probe was synthesized *in vitro* from pSPT19 AS1-S using T7 RiboMAX Large Scale RNA Production Systems (Promega, Madison, WI, USA). Briefly, approximately 1 µg of the plasmid was mixed with T7 polymerase and incubated in 37 °C for 4 hr. After transcription, the synthesized RNA was purified and biotin-labeled using Pierce RNA 3’ End Biotinylation Kit (Thermo Fisher Scientific). The obtained biotin-labeled RNA was used as an RNA probe. Similarly, an RNA probe that is the complement to the AS1-S sequence was synthesized from pSPT19 AS1-S using SP6 RiboMAX Large Scale RNA Production System (Promega) and used as a control probe. All procedures were performed in accordance with the manufacturer’s instructions.

Pull-down assays were performed using Pierce Magnetic RNA-Protein Pull-Down Kit (Thermo Fisher Scientific) in accordance with the manufacturer’s instructions. Briefly, 2 µg of the RNA probe was mixed with streptavidin magnetic beads and incubated 4 °C for 1 hr, followed by mixing with BL3.1 cell lysates prepared using RIPA buffer (Thermo Fisher Scientific). After incubation 4 °C for 1 hr, the beads were washed three times with RIPA buffer, and the RNA-protein complex was suspended using elution buffer. The samples were subjected to sodium dodecyl sulfate-polyacrylamide gel electrophoresis (SDS-PAGE) and stained using a Silver Stain MS Kit (FUJIFILM Wako, Tokyo, Japan). Protein bands specifically observed in the AS1-S RNA probe sample were cut out from the gel and subjected to LC-MS. LC-MS was performed by Japan Proteomics Co., Ltd. (https://www.jproteomics.com/).

### SDS-PAGE and western blotting

Samples were mixed with 4 × Laemmli sample buffer (BIO-RAD, Hercules, CA, USA) containing 200 mM dithiothreitol (DTT) and boiled for 5 min at 95 °C. The protein was separated by a 5–20% polyacrylamide gradient gel (ATTO, Tokyo, Japan) and transferred to a polyvinylidene difluoride (PVDF) membrane using iBlot 2 Dry Blotting System (Thermo Fisher Scientific). The membrane was then incubated in 5% skim milk (FUJIFILM Wako) in T-PBS buffer (PBS containing 0.05% Tween 20) at room temperature for 1 hr. After the blocking step, membranes were incubated with primary antibodies (rabbit anti-hnRNPM MAb (abcam: ab177957), mouse anti-BLV gp51 MAb (VMRD: BLV2), mouse anti-BLV p24 MAb (VMRD: BLV3), mouse anti-His tag MAb (MBL: D291-3), mouse anti-beta-tubulin MAb (Merck: 05-661)) in 5% skim milk in T-PBS buffer at room temperature for 1 hr. After a washing step, the membrane was incubated with the corresponding peroxidase-conjugated secondary antibodies (goat anti-mouse IgG MAb (abcam: ab6789), goat anti-rabbit IgG MAb (ZYMED Laboratories, Carlsbad, CA, USA: 82-6120)) in 5% skim milk in T-PBS buffer at room temperature for 1 hr. The detected proteins were visualized using the Super Signal West Dura Extended Duration Substrate (Thermo Fisher Scientific), and the image was obtained by imaging device FluorChem FC2 (ProteinSimple, Tokyo, Japan).

### RNA immunoprecipitation (RIP) assay

Magna RIP kit (Merck) was used to perform the RIP assay in accordance with the manufacturer’s instructions. Briefly, 2 µg of mouse anti-hnRNPM MAb (Santacruz: 5-RE36) and a control antibody (component of the kit) were mixed with magnetic beads and incubated at room temperature for 30 min. Subsequently, the beads were mixed with a BL3.1 cell lysate that was prepared using the lysis buffer included in the kit. After incubation at 4°C for 3 hr, beads were washed six times with the wash buffer included in the kit, and the obtained protein-RNA complexes were utilized in subsequent experiments.

### Conventional PCR and real-time PCR

For conventional reverse transcriptase (RT)-PCR, total RNA was extracted from cells using RNeasy Mini Kit (QIAGEN, Tokyo, Japan), followed by RT-PCR using PrimeScript One Step RT-PCR Kit Ver.2 (TaKaRa) with primers no. 7 and 8 (for detecting full-length of AS1-S) or 9 and 10 (for GAPDH). The PCR conditions were as follow; 50° C for 30 min, 94 °C for 2 min, followed by 35 cycles of 94 °C for 30 s, 60° C for 30 s, and 72 °C for 30 s.

RNAs encoding U6 and TYR were used as controls for nuclear and cytoplasmic RNA, respectively, and were quantified by real-time RT-PCR using One Step TB Green PrimeScript RT-PCR Kit II (Perfect Real Time) (TaKaRa) and QuantStudio 3 Real-Time PCR System (Thermo Fisher Scientific) with primers no. 11 and 12 or 13 and 14. AS1-S RNA in established MDBK CAG AS1-S and MDBK 3’LTR AS1-S cells were quantified in the same manner using primers no. 15 and 16. For quantification of AS1-S RNA in BL3.1 cells, strand-specific real-time RT-PCR was performed as previously reported (20) in order to delineate sense and anti-sense transcript levels. Briefly, RNA samples were reverse-transcribed using Prime Script RT Reagent kit (TaKaRa) in combination with an AS1-specific tagged primer (no. 17). The resultant reaction mixture was then 10-fold diluted and subjected to quantitative PCR using TB Green Premix Ex Taq II kit (TaKaRa) in combination with a tag primer (no. 18) and an AS1-specific reverse primer (no. 19). All PCR conditions were performed in accordance with the manufacturer’s instructions.

### Analysis of sub-cellular localization of RNAs

Nuclear or cytoplasmic RNA were separately extracted in accordance with a protocol provided from QIAGEN with slight modification (https://www.qiagen.com/us/resources/download.aspx?id=1f8ed6e7-2423-4cff-8013-0eb89ead5cb0&lang=en&ver=1: accessed 30 Oct. 2022). Briefly, cell pellets were lysed with RLN buffer (50 mM Tris-HCl, 140 mM NaCl, 1.5 mM MgCl_2_, 0.5% NP40), and nuclear and cytoplasmic fractions were separated by centrifugation, followed by RNA extraction using NucleoSpin RNA Plus (TaKaRa).

### RNA-seq and transcriptome analysis

For RNA-seq-based transcriptome analysis, total RNA was extracted from MDBK 3’LTR AS1-S, MDBK mock, and parental MDBK cells using NucleoSpin RNA Plus (TaKaRa) (n=2). Construction of an RNA-seq library and sequencing procedure were performed by Macrogen Japan Co., Ltd. (https://www.macrogen-japan.co.jp/); sequencing libraries were prepared using TruSeq stranded mRNA LT Sample Prep Kit (Illumina, San Diego, CA, USA), and next-generation sequencing analysis was conducted using NovaSeq 6000 (100-bp paired-end reads, approximately 4-Gbp total read bases per sample). The obtained data were trimmed using Trimmomatic (ver. 0.36) and then mapped to the reference genome (Bos taurus genome: ARS-UCD1.2) using HISAT2 (ver. 2.0.4), followed by transcript assembly and quantification using StringTie (ver. 2.1.1). After the assembly step, genes read counts of < 10 in at least one sample were removed. Principal component analysis (PCA) was performed using prcomp (Stats package in R ver. 4. 2. 1), and differential expression analysis was carried out using DESeq2 (ver. 1.36.0), followed by gene ontology analysis using clusterProfiler ((21), ver. 4.4.4). Differentially expressed genes (DEGs) were calculated between MDBK 3’LTR AS1-S (n=2) and MDBK mock and parental cells (n=4) to identify specific changes caused by the introduction of AS1-S. Significant DEGs were defined based on the following criteria: |fold change| ≥ 2, exactTest raw p-value < 0.05. For analysis of mRNA distribution, nuclear and cytoplasmic RNA were subjected to RNA-seq, followed by data manipulation as described above for transcriptome analysis.

### Analysis of alternative splicing using rMATS

Alternative splicing analysis was performed with rMATS software ((22), ver. 4.1.2) using bam files from the transcriptome analysis. To identify significant events in AS1-S expressing cells, splicing profiles from MDBK 3’LTR AS1-S cells (n=2) and MDBK mock and parental cells (n=4) were compared. The rMATS output showed total and differential alternative splicing events (ASEs); the following five event types were enumerated: skipped exons (SE), alternative 5’ splice sites (A5SS), alternative 3’ splice sites (A3SS), mutually exclusive exons (MXE), and retained introns (RI). Significantly different events were calculated using the differential percent spliced-in (PSI) between two groups; the result are presented as delta PSI values ranging from -1.0 to 1.0, and events with FDR < 0.1 were defined as significant. After rMATS analysis, the obtained gene list of significant differential ASEs was subjected to gene ontology analysis using clusterProfiler (ver. 4.4.4).

### RIP-seq analysis

RNA obtained from the RIP assay sample was purified using NucleoSpin RNA Plus (TaKaRa), followed by library construction and RNA-seq analysis. Construction of RNA-seq libraries and the sequencing procedure were performed by Macrogen Japan Co., Ltd. The sequence library was prepared using SMARTer Stranded RNA-Seq Kit (Illumina, San Diego, CA, USA), and next-generation sequencing analysis was conducted using NovaSeq 6000 (100-bp paired-end reads, approximately 4-Gbp total read bases per sample). The obtained data were filtered using prinseq (ver. 0.20.4) to remove abnormal reads with GC content < 20 and > 70. Subsequently, the obtained reads were trimmed using Trimmomatic (ver. 0.36) and mapped to the reference genome (Bos taurus genome: ARS-UCD1.2) using HISAT2 (ver. 2.0.4), followed by transcript assembly and quantification using StringTie (ver. 2.1.1). After assembly, genes with read counts of 0 in all samples were removed, and read counts in each gene were compared between RIP-assay samples from the anti-hnRNPM MAb and the control Ab in each cell line. Genes that meet the criteria described below were defined as hnRNPM-binding RNAs: read counts > 10, hnRNPM/control ratio > 2.0. The resultant hnRNPM-binding RNA lists were compared between MDBK 3’LTR AS1-S and MDBK mock cells, and genes that were observed in only MDBK 3’LTR AS1-S cells were subjected to gene ontology analysis using clusterProfiler (ver. 4.4.4).

### Appendices

The primers used in this study are listed in Table S1 in the supplemental material. The FASTQ data in this study were disclosed in the DDBJ database as submission no. DRR444704–444720 and DRR446101. All supplemental information are shown in the supplemental material.

## Acknowledgments

This study was supported by JSPS KAKENHI Grant-in-Aid for Early-Career Scientists (20K15687) (KA) and the Ito Foundation Research Grant from FY2020 to FY2022 (KA).

Conceptualization, K.A.; Data curation, K.A., A.N., Y.M.; Formal analysis, K.A.; Funding acquisition, K.A.; Investigation, K.A.; Methodology, K.A., A.N., Y.M.; Project administration, K.A.; Resources, K.A., A.N., Y.M.; Software, K.A.; Supervision, K.A.; Validation. K.A.; Visualization, K.A.; Writing – original draft, K.A.; Writing – review & editing, K.A., A.N., Y.M.

We thank the members of our laboratory for their assistance and Forte (https://www.fortescience.com/) for the English language review.

The authors declare that they have no competing interests.

## SUPPLEMENTAL MATERIAL

SUPPLEMENTAL FILE 1 (supplemental figures)

SUPPLEMENTAL FILE 2 (supplemental tables)

---

**Fig. S1.**
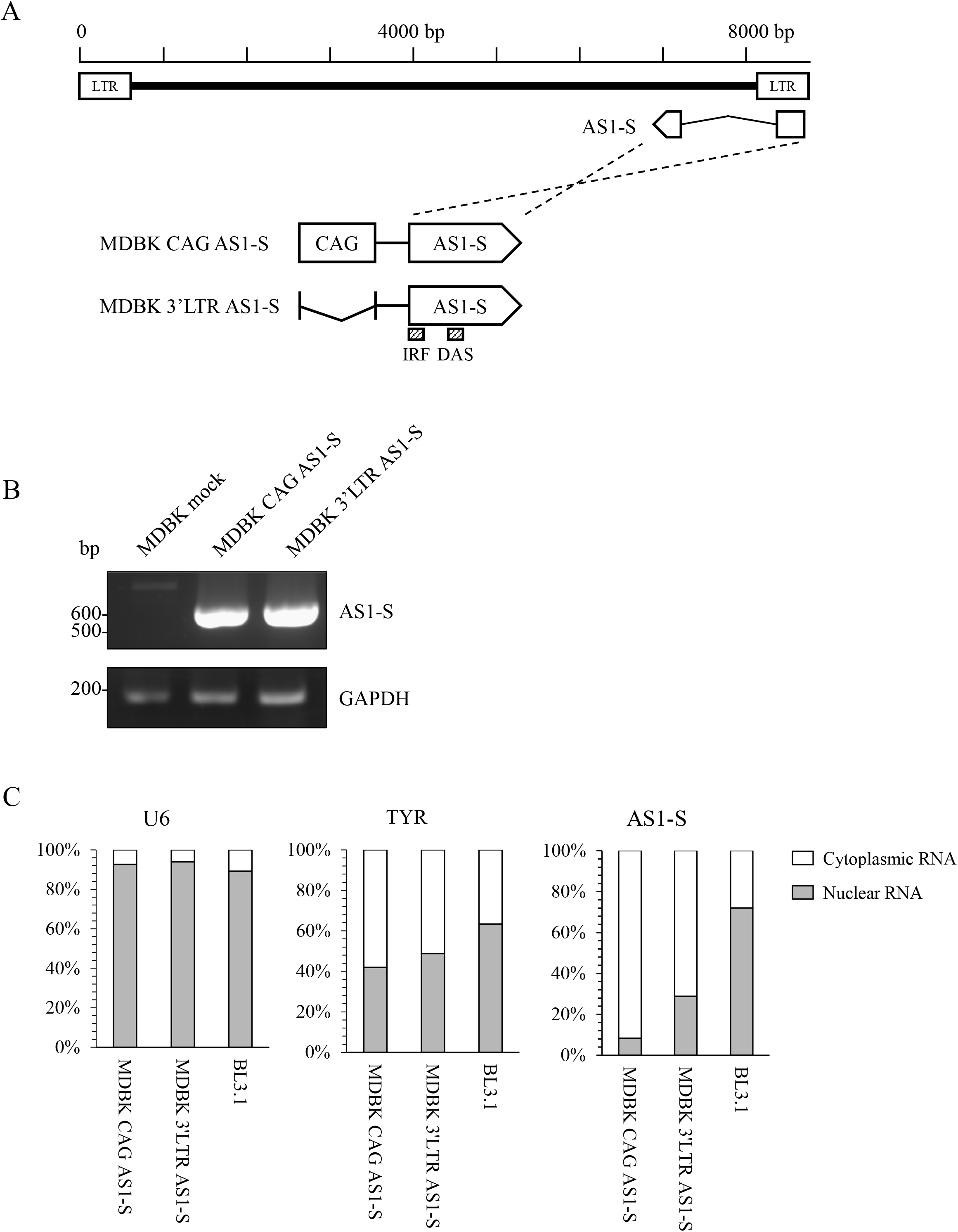
(A) Schematic diagrams of the AS1-S expression plasmids. CAG: CAG promoter, AS1-S: cDNA sequence of BLV AS1-S. IRF (interferon regulatory factor) and DAS (downstream activator sequence) indicate regulatory sequences reported in Durkin et al. (11). (B) Results of RT-PCR amplification of whole AS1-S RNA and the housekeeping gene *GAPDH*. Total RNA was extracted from transfected cells and subsequently subjected to conventional RT-PCR. (C) Sub-cellular localization of U6, TYR and AS1-S RNAs in transfected cells. Nuclear (grey) and cytoplasmic (white) RNA were separately extracted from the cells, followed by real-time RT-PCR. The cytoplasm/nucleus ratio for each RNA was calculated (n=3) and shown as percentages.

**Fig. S2.**
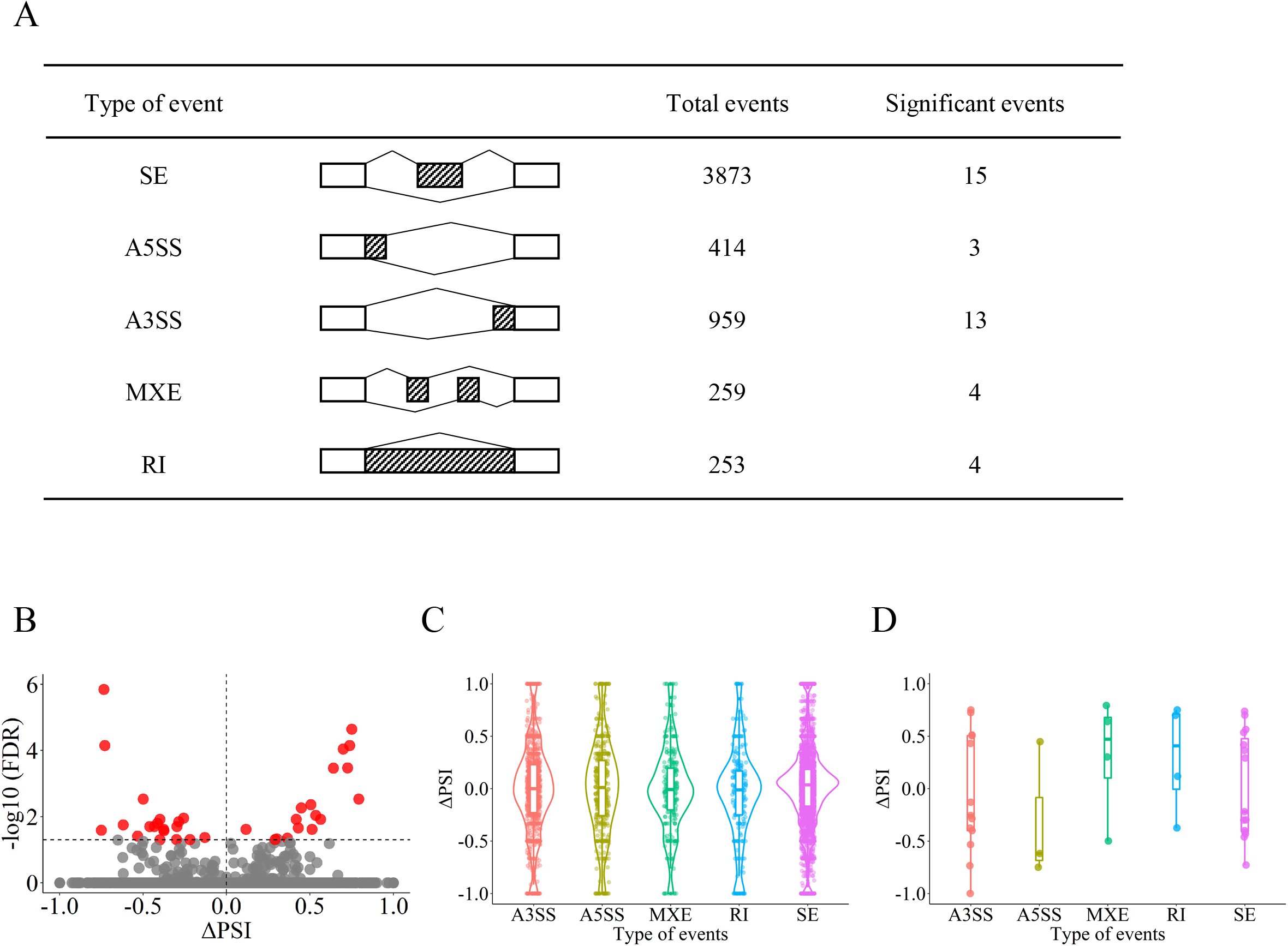
(A) Summary of the alternative splicing analysis using rMATS. The diagrams show the five splicing patterns: skipped exon (SE), alternative 5’ splice site (A5SS), alternative 3’ splice site (A3SS), mutually exclusive exons (MXE), and retained intron (RI). (B) Volcano plot of the splicing analysis. The X and Y axes indicate the delta PSI and FDR, respectively. Significant events are shown as red dots. (C, D) Violin plots of total (C) and significant (D) events identified by alternative splicing analysis using rMATs. MDBK 3’LTR AS1-S cells (n=2) and MDBK mock and parental cells (n=4) were compared, and events with FDR < 0.1 were defined as significant. The Y axis indicates the delta PSI between the two groups.

**Fig. S3.**
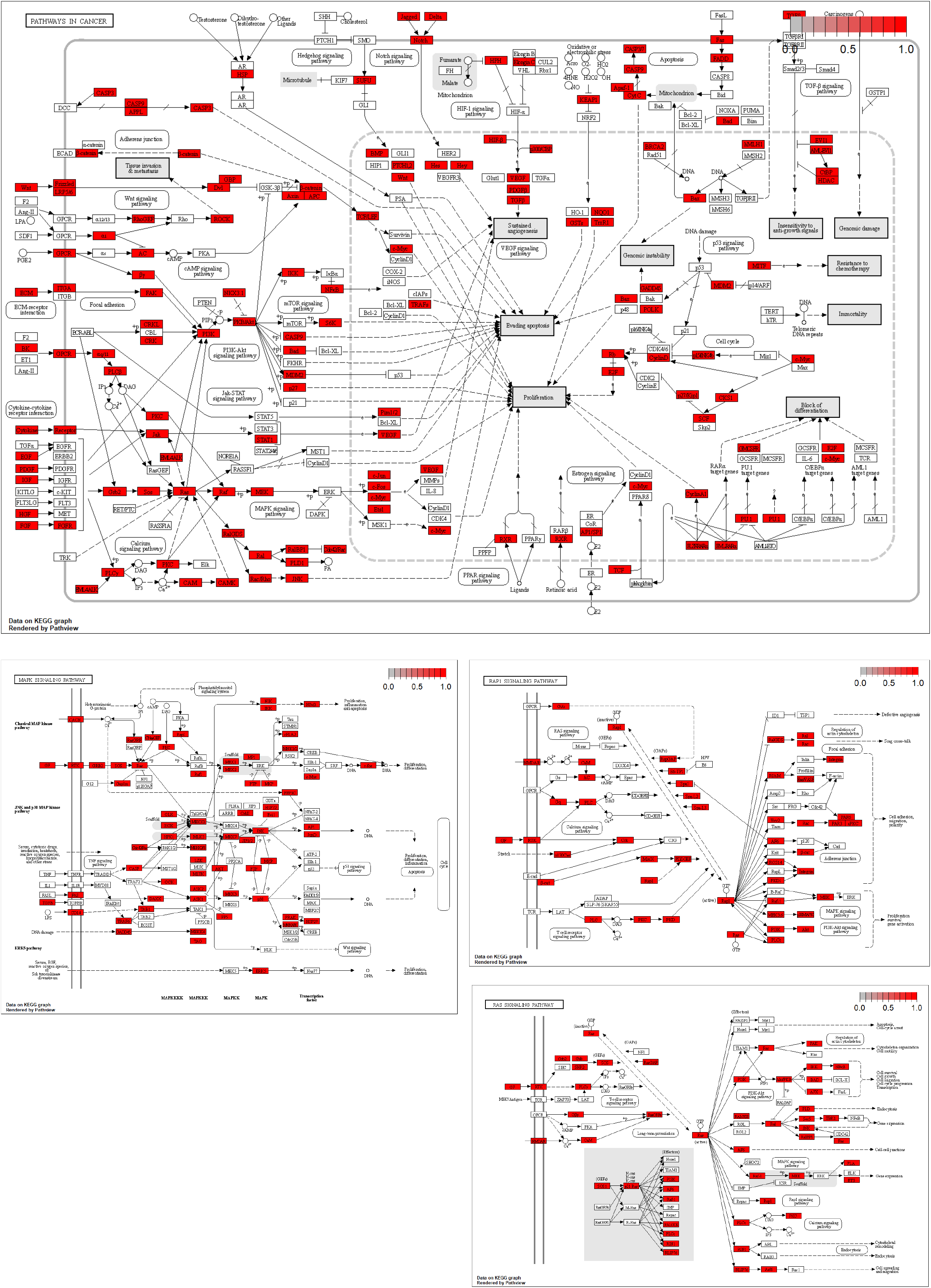
Diagrams of the KEGG pathway “bta05200 (Pathways in cancer)”, “bta04010 (MAPK signaling pathway)”, “bta04014 (Ras signaling pathway)”, and “bta04015 (Rap1 signaling pathway)”. Red genes indicate hnRNPM-binding RNAs detected in only MDBK 3’LTR AS1-S cells identified by RIP-seq.

**Fig. S4.**
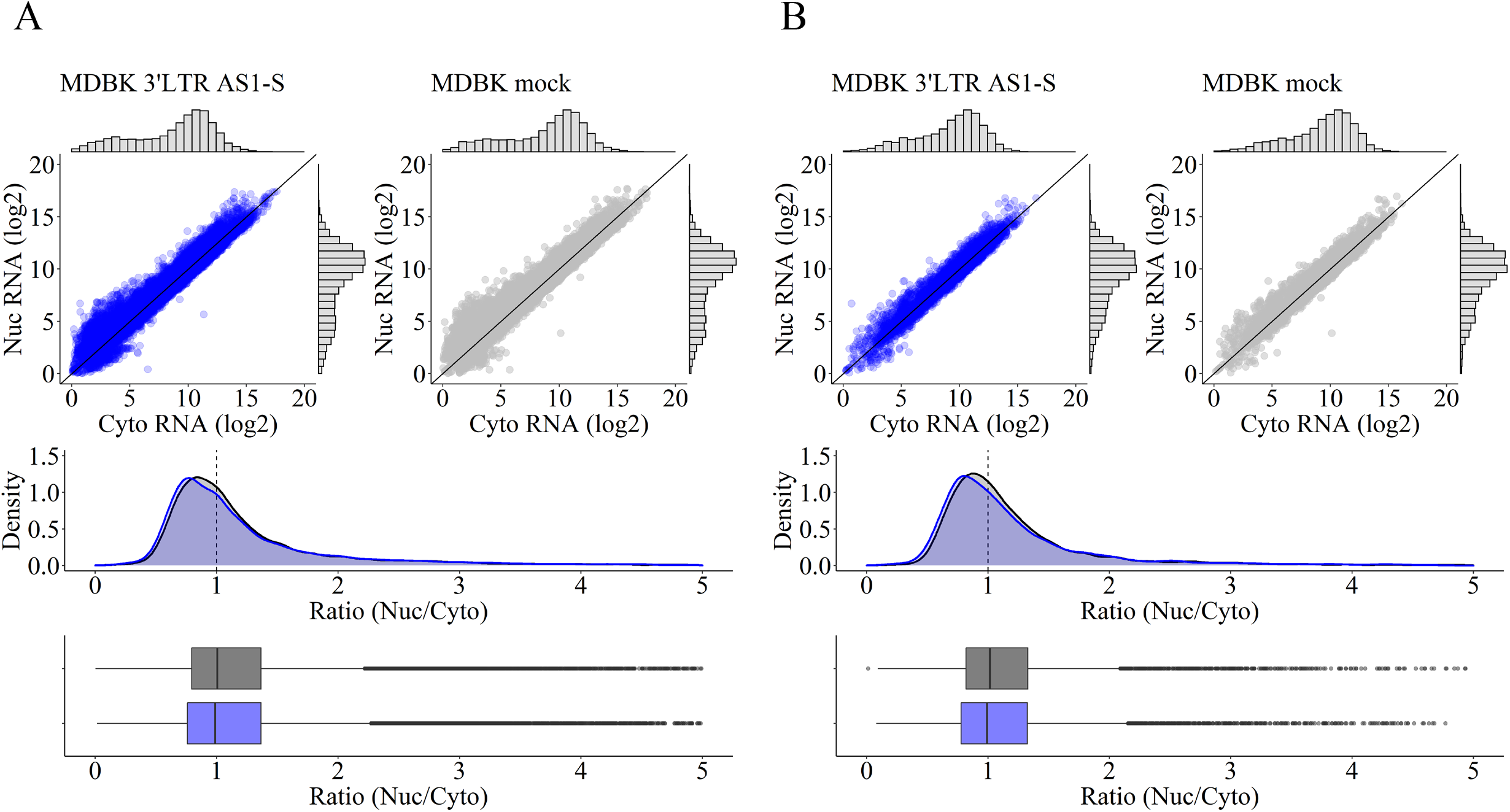
(A) Scatter plots showing read counts of nuclear (Nuc) and cytoplasmic (Cyto) RNA extracted from MDBK 3’LTR AS1-S and MDBK mock cells. Read counts were normalized using DEseq2 prior to being used for analysis. The X and Y axes show read counts of cytoplasmic (Cyto) and nuclear (Nuc) RNA, respectively. The bottom graph shows the density plot and box plot of the nuclear/cytoplasmic RNA ratio in MDBK 3’LTR AS1-S (blue) and MDBK mock (black) cells. The X axis shows the calculated ratio of nuclear and cytoplasmic RNA. The vertical lines indicate a Nuc/Cyto RNA ratio = 1.0. (B) mRNAs of the 4607 genes in Fig. 4C were extracted from the result shown in panel (A).

**Fig. S5.**
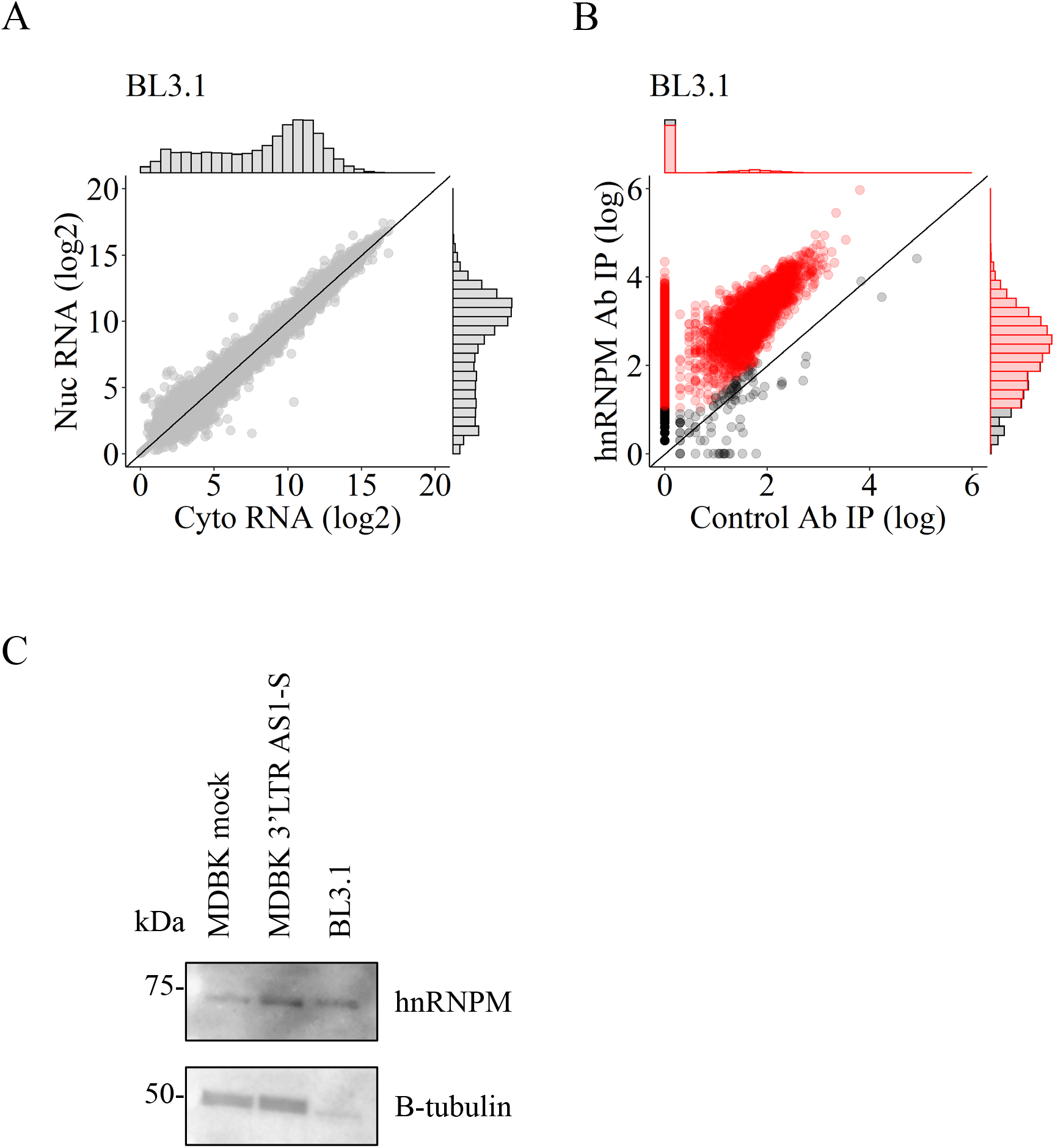
(A) Scatter plot showing read counts of nuclear (Nuc) and cytoplasmic (Cyto) RNA extracted from BL3.1 cells. Read counts were normalized using DEseq2 prior to being used for analysis. The X and Y axes show read counts of cytoplasmic (Cyto) and nuclear (Nuc) RNA, respectively. (B) Scatter plot showing sequencing reads obtained by RNA immunoprecipitation using BL3.1 cells. The X and Y axes show read counts obtained by RIP-seq from control MAb and anti-hnRNPM MAb, respectively. Genes that meet the following criteria were defined as hnRNPM-binding RNAs and are shown as red dots: read counts > 10, hnRNPM/control ratio > 2.0. (C) Western blot analysis comparing the expression levels of hnRNPM in BL3.1, MDBK 3’LTR AS1-S and MDBK mock cells. 1 × 10^6^ cells were subjected to the analysis.

## References

1. Virgin HW, Wherry EJ, Ahmed R. 2009. Redefining Chronic Viral Infection. Cell 138:30–50.

2. Grundhoff A, Sullivan CS. 2011. Virus-encoded microRNAs. Virology 411:325–343.

3. Pijlman GP, Funk A, Kondratieva N, Leung J, Torres S, van der Aa L, Liu WJ, Palmenberg AC, Shi P-Y, Hall RA, Khromykh AA. 2008. A Highly Structured, Nuclease-Resistant, Noncoding RNA Produced by Flaviviruses Is Required for Pathogenicity. Cell Host & Microbe 4:579–591.

4. Lee N, Yario TA, Gao JS, Steitz JA. 2016. EBV noncoding RNA EBER2 interacts with host RNA-binding proteins to regulate viral gene expression. Proc Natl Acad Sci USA 113:3221–3226.

5. Satou Y, Yasunaga J -i., Yoshida M, Matsuoka M. 2006. HTLV-I basic leucine zipper factor gene mRNA supports proliferation of adult T cell leukemia cells. Proceedings of the National Academy of Sciences 103:720–725.

6. Fabian MR, Sonenberg N, Filipowicz W. 2010. Regulation of mRNA Translation and Stability by microRNAs. Annu Rev Biochem 79:351–379.

7. Rinn JL, Chang HY. 2012. Genome Regulation by Long Noncoding RNAs. Annu Rev Biochem 81:145–166.

8. Kincaid RP, Burke JM, Sullivan CS. 2012. RNA virus microRNA that mimics a B-cell oncomiR. Proceedings of the National Academy of Sciences 109:3077–3082.

9. Kincaid RP, Chen Y, Cox JE, Rethwilm A, Sullivan CS. 2014. Noncanonical MicroRNA (miRNA) Biogenesis Gives Rise to Retroviral Mimics of Lymphoproliferative and Immunosuppressive Host miRNAs. mBio 5.

10. Rosewick N, Momont M, Durkin K, Takeda H, Caiment F, Cleuter Y, Vernin C, Mortreux F, Wattel E, Burny A, Georges M, Van den Broeke A. 2013. Deep sequencing reveals abundant noncanonical retroviral microRNAs in B-cell leukemia/lymphoma. Proceedings of the National Academy of Sciences 110:2306– 2311.

11. Durkin K, Rosewick N, Artesi M, Hahaut V, Griebel P, Arsic N, Burny A, Georges M, Van den Broeke A. 2016. Characterization of novel Bovine Leukemia Virus (BLV) antisense transcripts by deep sequencing reveals constitutive expression in tumors and transcriptional interaction with viral microRNAs. Retrovirology 13:33.

12. Gillet N, Florins A, Boxus M, Burteau C, Nigro A, Vandermeers F, Balon H, Bouzar A-B, Defoiche J, Burny A, Reichert M, Kettmann R, Willems L. 2007. Mechanisms of leukemogenesis induced by bovine leukemia virus: prospects for novel anti-retroviral therapies in human. Retrovirology 4:18.

13. Gillet NA, Gutiérrez G, Rodriguez SM, de Brogniez A, Renotte N, Alvarez I, Trono K, Willems L. 2013. Massive Depletion of Bovine Leukemia Virus Proviral Clones Located in Genomic Transcriptionally Active Sites during Primary Infection. PLoS Pathog 9:e1003687.

14. Andoh K, Kimura K, Nishimori A, Hatama S. 2020. Development of an in situ hybridization assay using an AS1 probe for detection of bovine leukemia virus in BLV-induced lymphoma tissues. Arch Virol 165:2869–2876.

15. Gillet NA, Hamaidia M, de Brogniez A, Gutiérrez G, Renotte N, Reichert M, Trono K, Willems L. 2016. Bovine Leukemia Virus Small Noncoding RNAs Are Functional Elements That Regulate Replication and Contribute to Oncogenesis In Vivo. PLoS Pathog 12:e1005588.

16. Safari R, Jacques J-R, Brostaux Y, Willems L. 2020. Ablation of non-coding RNAs affects bovine leukemia virus B lymphocyte proliferation and abrogates oncogenesis. PLoS Pathog 16:e1008502.

17. Andoh K, Nishimori A, Sakumoto R, Hayashi K-G, Hatama S. 2020. The chemokines CCL2 and CXCL10 produced by bovine endometrial epithelial cells induce migration of bovine B lymphocytes, contributing to transuterine transmission of BLV infection. Veterinary Microbiology 242:108598.

18. Nishimori A, Andoh K, Matsuura Y, Kumagai A, Hatama S. 2021. Establishment of a simplified inverse polymerase chain reaction method for diagnosis of enzootic bovine leukosis. Arch Virol 166:841–851.

19. Ma Q, Li L, Tang Y, Fu Q, Liu S, Hu S, Qiao J, Chen C, Ni W. 2017. Analyses of long non-coding RNAs and mRNA profiling through RNA sequencing of MDBK cells at different stages of bovine viral diarrhea virus infection. Research in Veterinary Science 115:508–516.

20. Andoh K, Akagami M, Nishimori A, Matsuura Y, Kumagai A, Hatama S. 2021. Novel single nucleotide polymorphisms in the bovine leukemia virus genome are associated with proviral load and affect the expression profile of viral non-coding transcripts. Veterinary Microbiology 261:109200.

21. Yu G, Wang L-G, Han Y, He Q-Y. 2012. clusterProfiler: an R Package for Comparing Biological Themes Among Gene Clusters. OMICS: A Journal of Integrative Biology 16:284–287.

22. Shen S, Park JW, Lu Z, Lin L, Henry MD, Wu YN, Zhou Q, Xing Y. 2014. rMATS: Robust and flexible detection of differential alternative splicing from replicate RNA-Seq data. Proc Natl Acad Sci USA 111:E5593–E5601.

23. Datar KV, Dreyfuss G, Swanson MS. 1993. The human hnRNP M proteins: identification of a methionine/arginine-rich repeat motif in ribonucleoproteins. Nucl Acids Res 21:439–446.

24. Gattoni R. 1996. The human hnRNP-M proteins: structure and relation with early heat shock-induced splicing arrest and chromosome mapping. Nucleic Acids Research 24:2535–2542.

25. Piñol-Roma S, Dreyfuss G. 1993. hnRNP proteins:Localization and transport between the nucleus and the cytoplasm. Trends in Cell Biology 3:151–155.

26. Neriec N, Percipalle P. 2018. Sorting mRNA Molecules for Cytoplasmic Transport and Localization. Front Genet 9:510.

27. Unfried JP, Ulitsky I. 2022. Substoichiometric action of long noncoding RNAs. Nat Cell Biol 24:608–615.

28. Murakami H, Todaka H, Uchiyama J, Sato R, Sogawa K, Sakaguchi M, Tsukamoto K. 2019. A point mutation to the long terminal repeat of bovine leukemia virus related to viral productivity and transmissibility. Virology 537:45–52.

29. Tajima S, Takahashi M, Takeshima S, Konnai S, Yin SA, Watarai S, Tanaka Y, Onuma M, Okada K, Aida Y. 2003. A Mutant Form of the Tax Protein of Bovine Leukemia Virus (BLV), with Enhanced Transactivation Activity, Increases Expression and Propagation of BLV In Vitro but Not In Vivo. J Virol 77:1894–1903.

30. Aida Y, Murakami H, Takahashi M, Takeshima S-N. 2013. Mechanisms of pathogenesis induced by bovine leukemia virus as a model for human T-cell leukemia virus. Front Microbiol 4.

31. Lo C-W, Borjigin L, Saito S, Fukunaga K, Saitou E, Okazaki K, Mizutani T, Wada S, Takeshima S, Aida Y. 2020. BoLA-DRB3 Polymorphism is Associated with Differential Susceptibility to Bovine Leukemia Virus-Induced Lymphoma and Proviral Load. Viruses 12:352.

32. Lo C-W, Takeshima S, Okada K, Saitou E, Fujita T, Matsumoto Y, Wada S, Inoko H, Aida Y. 2021. Association of Bovine Leukemia Virus-Induced Lymphoma with BoLA-DRB3 Polymorphisms at DNA, Amino Acid, and Binding Pocket Property Levels. Pathogens 10:437.

33. Sajiki Y, Konnai S, Okagawa T, Nishimori A, Maekawa N, Goto S, Watari K, Minato E, Kobayashi A, Kohara J, Yamada S, Kaneko MK, Kato Y, Takahashi H, Terasaki N, Takeda A, Yamamoto K, Toda M, Suzuki Y, Murata S, Ohashi K. 2019. Prostaglandin E _2_ –Induced Immune Exhaustion and Enhancement of Antiviral Effects by Anti–PD-L1 Antibody Combined with COX-2 Inhibitor in Bovine Leukemia Virus Infection. JI 203:1313–1324.

34. Konnai S, Murata S, Ohashi K. 2017. Immune exhaustion during chronic infections in cattle. The Journal of Veterinary Medical Science 79:1–5.

35. Rosewick N, Durkin K, Artesi M, Marçais A, Hahaut V, Griebel P, Arsic N, Avettand-Fenoel V, Burny A, Charlier C, Hermine O, Georges M, Van den Broeke A. 2017. Cis-perturbation of cancer drivers by the HTLV-1/BLV proviruses is an early determinant of leukemogenesis. Nat Commun 8:15264.

36. Cao P, Luo W-W, Li C, Tong Z, Zheng Z-Q, Zhou L, Xiong Y, Li S. 2019. The heterogeneous nuclear ribonucleoprotein hnRNPM inhibits RNA virus-triggered innate immunity by antagonizing RNA sensing of RIG-I-like receptors. PLoS Pathog 15:e1007983.

37. West KO, Scott HM, Torres-Odio S, West AP, Patrick KL, Watson RO. 2019. The Splicing Factor hnRNP M Is a Critical Regulator of Innate Immune Gene Expression in Macrophages. Cell Reports 29:1594–1609.e5.

38. Passacantilli I, Frisone P, De Paola E, Fidaleo M, Paronetto MP. 2017. hnRNPM guides an alternative splicing program in response to inhibition of the PI3K/AKT/mTOR pathway in Ewing sarcoma cells. Nucleic Acids Research 45:12270–12284.

39. Xu Y, Gao XD, Lee J-H, Huang H, Tan H, Ahn J, Reinke LM, Peter ME, Feng Y, Gius D, Siziopikou KP, Peng J, Xiao X, Cheng C. 2014. Cell type-restricted activity of hnRNPM promotes breast cancer metastasis via regulating alternative splicing. Genes Dev 28:1191–1203.

40. Shiokawa M, Miura R, Okubo A, Hagita Y, Yoshimura I, Aoki H. 2021. Bovine endometrium-derived cultured cells are suitable for lipofection. Sci Rep 11:16207.

41. Morizako N, Butlertanaka EP, Tanaka YL, Shibata H, Okabayashi T, Mekata H, Saito A. 2022. Generation of a bovine cell line for gene engineering using an HIV-1-based lentiviral vector. Sci Rep 12:16952.

42. Toyoda K, Matsuoka M. 2022. Functional and Pathogenic Roles of Retroviral Antisense Transcripts. Front Immunol 13:875211.

43. Matsuoka M, Mesnard J-M. 2020. HTLV-1 bZIP factor: the key viral gene for pathogenesis. Retrovirology 17:2.

44. Mitobe Y, Yasunaga J, Furuta R, Matsuoka M. 2015. HTLV-1 bZIP Factor RNA and Protein Impart Distinct Functions on T-cell Proliferation and Survival. Cancer Research 75:4143–4152.

45. Gazon H, Chauhan PS, Porquet F, Hoffmann GB, Accolla R, Willems L. 2020. Epigenetic silencing of HTLV-1 expression by the HBZ RNA through interference with the basal transcription machinery. Blood Advances 4:5574–5579.

